# RECTA: Regulon Identification Based on Comparative Genomics and Transcriptomics Analysis

**DOI:** 10.1101/261453

**Authors:** Xin Chen, Anjun Ma, Adam McDermaid, Hanyuan Zhang, Chao Liu, Huansheng Cao, Qin Ma

## Abstract

Regulons, which serve as co-regulated gene groups contributing to the transcriptional regulation of microbial genomes, have the potential to aid in understanding of underlying regulatory mechanisms. In this study, we designed a novel computational pipeline, RECTA, for regulon prediction related to the gene regulatory network under certain conditions. To demonstrate the effectiveness of this tool, we implemented RECTA on *Lactococcus lactis* MG1363 data to elucidate acid-response regulons. *Lactococcus lactis* is one of the most important Gram-positive lactic acid-producing bacteria, widely used in food industry and has been proved to have advantages in oral delivery of drug and vaccine. The pipeline carries out differential gene expression, gene co-expression analysis, *cis*-regulatory motif finding, and comparative genomics to predict and validate regulons related to acid stress response. A total of 51 regulons were identified, 14 of which have computational-verified significance. Among these 14 regulons, five of them were computationally predicted to be connected with acid stress response with (i) known transcriptional factors in MEME suite database successfully mapped in *Lactococcus lactis* MG1363; and (ii) differentially expressed genes between pH values of 6.5 (control) and 5.1 (treatment). Validated by 36 literature confirmed acid stress response related proteins and genes, 33 genes in *Lactococcus lactis* MG1363 were found having orthologous genes using BLAST, associated to six regulons. An acid response related regulatory network was constructed, involving two trans-membrane proteins, eight regulons (*llrA, llrC, hllA, ccpA*, NHP6A, *rcfB*, regulons #8 and #39), nine functional modules, and 33 genes with orthologous genes known to be associated to acid stress. Our RECTA pipeline provides an effective way to construct a reliable gene regulatory network through regulon elucidation. The predicted response pathways could serve as promising candidates for better acid tolerance engineering in *Lactococcus lactis*. RECTA has strong application power and can be effectively applied to other bacterial genomes where the elucidation of the transcriptional regulation network is needed.

## INTRODUCTION

Genomic and transcriptomic analyses have been widely used for elucidating gene regulatory network (GRN) hierarchies and offering insight into the coordination of response capabilities in microorganisms (Arnoldini et al. 2012, Carvalho et al. 2013, Levine et al. 2013, Locke et al. 2011). One way to study the mechanism of transcriptional regulation in microbe genomics is regulon prediction. A regulon is a group of co-regulated operons, which contains single or multiple consecutive genes along the genome (Cao, Ma, et al. 2017, Mao et al. 2015, Zhou et al. 2014). Genes in the same operon are controlled by the same promoter and are co-regulated by one or a set of TFs (Jacob et al. 1960). The elucidation of regulons can improve the identification of transcriptional genes, and thus, reliably predict the gene transcription regulation networks (Liu, Zhou, et al. 2016).

There are three ways for regulon prediction: (i) Predicting new operons for a known regulon (Kumka and Bauer 2015, Tan et al. 2001). This method combines motif profiling with a comparative genomic strategy to search for related regulon members and carries out systematical gene regulation study. (ii) Integrating *cis*-regulatory motif (*motif* for short) comparison and clustering to find significantly enriched motif candidates (Gupta et al. 2007, Ma et al. 2013). The candidate motifs are then assembled into regulons. (iii) Performing *ab initio* novel regulon inference using *de novo* motif finding strategy (Novichkov et al. 2010). This approach uses a phylogenetic footprinting technique which mostly relies on reference verification (Blanchette et al. 2002, Katara et al. 2012, Liu, Zhang, et al. 2016) and can perform a horizontal sequential comparison to predict regulons in target organisms by searching known functionally-related regulons or TFs from other relevant species. One algorithm for phylogenetic footprinting analysis called Motif Prediction by Phylogenetic Footprinting (MP3) has been used for regulon prediction in *E. coli* (Liu, Zhang, Zhou, Li, Fennell, Wang, Kang, Liu and Ma 2016). MP3 was then integrated into the DMINDA web server along with other algorithms, such as the Database of Prokaryotic Operons 2.0 (DOOR2) (Cao, Ma, Chen and Xu 2017, Mao et al. 2014), BOttleneck BROken (BoBro) (Li, Ma, et al. 2011) and BoBro-based motif Comparison (BBC) (Ma, Liu, Zhou, Yin, Li and Xu 2013), to construct a complete pipeline for regulon prediction. In the latest research, a newly developed pipeline called Single-cell Regulatory Network Inference and Clustering (SCENIC) combines motif finding from co-expression gene modules (CEMs) with regulon prediction for single-cell clustering and analysis (Aibar et al. 2017). Such method builds up a way of regulon application in single-cell and metagenomic research. Nevertheless, without a suitable regulon database, researchers need to build up the library first through operon identification, CEM analysis, motif prediction and comparison (Jensen et al. 2005). Here, we reported an integrated computational framework of <u>re</u>gulon identification based on <u>c</u>omparative genomics and transcriptomics <u>a</u>nalysis (**RECTA**) to elucidate the GRN responses in microbes under specific conditions. To better elucidate the methodology of RECTA, we built a regulatory network responding to the acid stress in *Lactococcus lactis* (*L. lactis*) species.

*Lactococcus lactis* (*L. lactis*) is one of the mesophilic Gram-positive lactic acid-producing bacteria. It is has been widely applied in dairy fermentations, such as cheese and milk product (Wegmann et al. 2007). Several studies have provided evidence of its essential roles in wrapping and delivering proteins or vaccinations for immune treatment, such as diabetes (Ma, Liu, et al. 2014), malaria (Ramasamy et al. 2006), tumors (Bermudez-Humaran et al. 2005, Zhang et al. 2016) and infections (Hanniffy et al. 2007). Holding the advantage of higher acid tolerance to protect vectors from resolving during delivery inside of the animal body, *L. lactis* has more potential and safety in oral drug development (Hols et al. 1999). Moreover, it has been found that *L. lactis*, along with some *Lactobacillus, Bifidobacterium* and other gut microbiota, were associated with obesity (Million et al. 2012). Such studies lead to the possibility and availability of *L. lactis* in metagenomic studies to investigate the effect of microbial interaction between *L. lactis* and other species in the human body. It is now well established that *Lactococcus* have evolved stress-sensing systems, which enable them to tolerate harsh environmental conditions (Carvalho, Turner, Fonseca, Solopova, Catarino, Kuipers, Voit, Neves and Santos 2013, Hutkins and Nannen 1993, van de Guchte et al. 2002).

Among the harsh environmental conditions that microorganisms confront, acid stress is known to change the level of the alarmones (guanosine tetraphosphate and guanosine pentaphosphate, collectively referred to as (p)ppGpp) (Hauryliuk et al. 2015) and leads to a stringent response to cellular regulation (Rallu et al. 2000). The reason that bacteria maintain the protection mechanism against acid stress is to withstand the deleterious effects caused by the harmful high level of protons in the exposed environment. Many mechanisms or genes related to the acid stress response (ASR) have been identified. Proton–pumping activity, the direct regulator to acid stress response, controls the intracellular pH level by pumping extra protons out of the cell (Koebmann et al. 2000, Lund et al. 2014), and the increase of alkaline compound levels also counters the acidification found in streptococci (Shabayek and Spellerberg 2017). Acid damage repair of cells by chaperones or proteases, such as GroES, GroEL, GrpE, HrcA, DnaK, DnaJ and Clp (Frees et al. 2003, Jayaraman et al. 1997), and hdeA/B and Hsp31 in *Escherichia coli* (*E. coli*) (Kern et al. 2007, Mujacic and Baneyx 2007), the arginine deiminase system (ADI) (Budin-Verneuil et al. 2006, Ryan et al. 2009, Sun et al. 2012, Zuniga et al. 2002) and glutamate decarboxylases (GAD) pathways, etc. (Hoskins et al. 2001, Nomura et al. 1999, Sanders et al. 1998) have been proven to be associated with the acid response. Additionally, transcriptional regulators, sigma factors, and two-component signal transduction system (TCSs) have also been demonstrated to be responsible for ASR by modifying gene expression (Cotter and Hill 2003). These genes or pathways suggest low pH has widespread adverse effects on cell functions and inflicts response at genomic, metabolic, and macromolecular levels. To better understand the mechanism that controls the acid tolerance and response to the acid stress in *L. lactis*, we considered MG1363, a strain extensively studied for acid resistance, to carry out computational analyses (Carvalho, Turner, Fonseca, Solopova, Catarino, Kuipers, Voit, Neves and Santos 2013, Hartke et al. 1996, Linares et al. 2010, Sanders et al. 1999). Nevertheless, to adequately describe the transcriptional state and gene regulation responsible for ASR in *L. lactis*, a GRN integrating all individual pathways is needed.

The experiment was conducted by six steps and the general framework is showcased in Figure 1. (i) MG1363 co-expression gene modules (CEMs) and differentially expressed genes (DEG) were generated from microarray data by hcluster package (Antoine Lucas 2006) and Wilcoxon test (Bauer 1972) in R, respectively. MG1363 operons were predicted from genome sequence using DOOR2 webserver and assigned into each CEM; (ii) For each CEM, the 300bp upstream to the promoter was extracted and the sequences were used to find motifs using DMINDA2.0; (iii) The top five significant motifs in each CEM were reassembled by their similarity comparison and clustering to predict regulons; (iv) The motifs were compared to known transcription factor binding sites (TFBSs) in the MEME suite, and the TFs corresponding to these TFBSs were mapped to MG1363 using BLAST. Only regulons with DEGs and mapped TF were kept as ASR-related regulons; (v) Experimentally identified ASR-related genes in other organisms were mapped to MG1363 using BLAST and allocated to corresponding regulons for further verification; and (vi) The relationship between regulons and functional gene modules were established to elucidate the overall ASR mechanism in MG1363.

**Figure 1:**
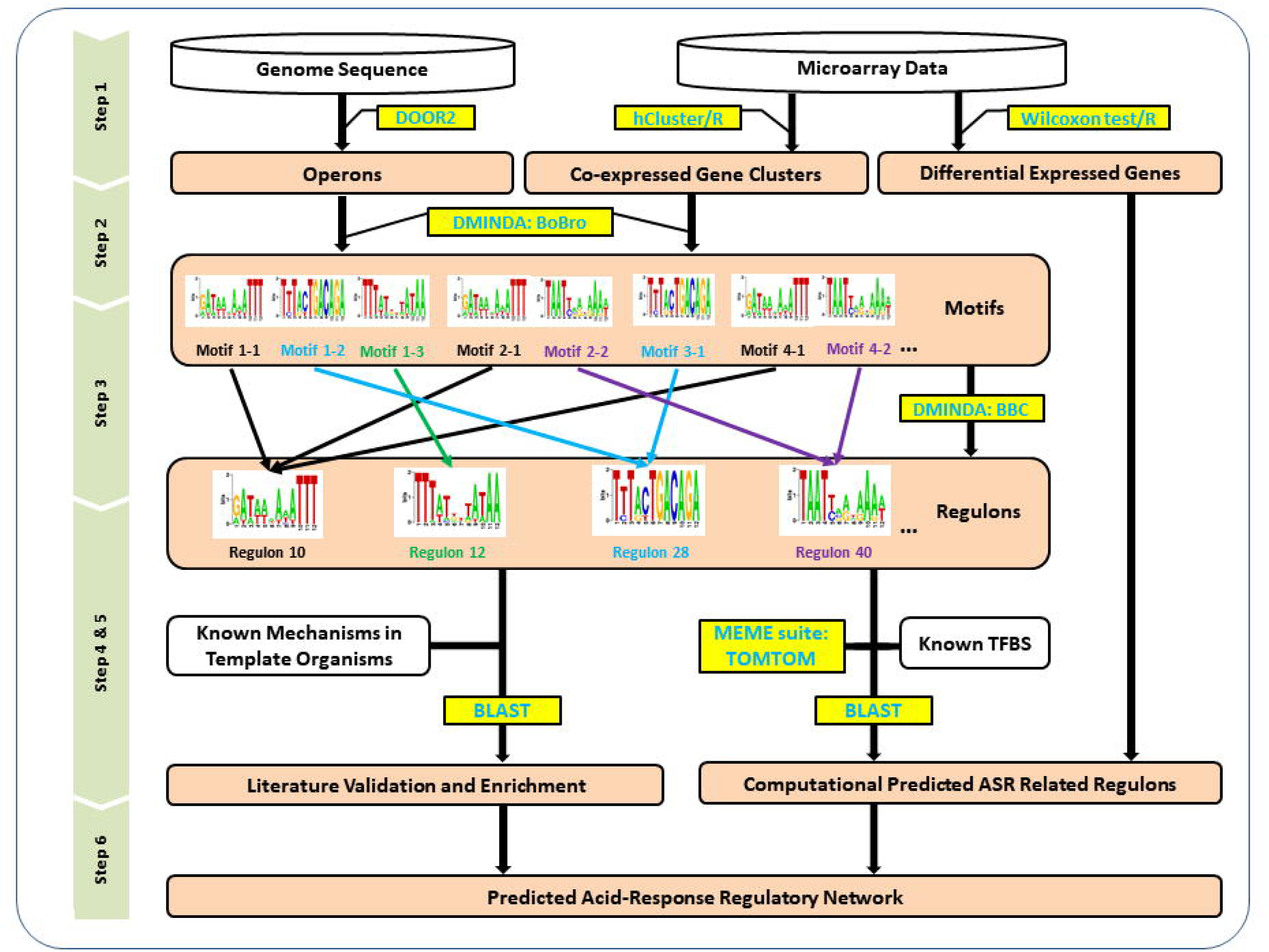
The flowchart of constructing global ASR transcriptional network in MG1363. Step 1: microarray data was used to generate co-expressed gene clustersand DEGs, and MG1363 genome sequence was used to find operons. Step 2: a motif finding progress was carried out to identify all statistically significant motifs in each of the CEMs. Step 3: a regulon finding procedure was designed to identify all the possible regulon candidates encoded in the genome based on motif comparison and clustering. Step 4: the motifs of each of these regulons were compared to known TFBSs, and DGE analysis between low pH condition and normal condition was used to figure out the ASR-related regulons. Step 5: regulon validation based on literature information verified the significant putative regulons and expanded the results to some insufficiently significant regulons. Step 6: the ASR-related GRN in MG1363 was predicted and described with eight regulons, nine functional modules, and 33 genes. The combination of the above information forms a genome-scale regulatory network constructed for ASR.

As a result, 14 regulons are identified, literature verified or putative, to be connected to ASR. Eight regulons, related to nine functional modules and 33 associated genes, are considered as the essential elements in acid resistance in MG1363. This proposed computational pipeline and the above results significantly expand the current understanding of the ASR system, providing a new method to predict systematic regulatory network based on regulon clustering.

## METHODS

### Data Acquisitio

The *L. lactic* MG1363 genome sequence was downloaded from NCBI (GenBank accession number: AM406671). The microarray dataset contains eight samples under different acid stress conditions for MG1363 was downloaded from the Gene Expression Omnibus (GEO) Database (Series number: GSE47012). The data has been treated with LOWESS normalization by the provider. The details on cell culture preparation and data processing can be found in the previous study (Carvalho, Turner, Fonseca, Solopova, Catarino, Kuipers, Voit, Neves and Santos 2013). This dataset has all bacteria grown in basic conditions: a 2-liter fermenter in chemically defined medium containing 1% (w/v) glucose at 30°C. The control and treatment samples were grown at a pH of 6.5 and 5.1, respectively.

Several TFBS databases integrated in the MEME suite, including DPINTERACT (*E. coli*) (http://arep.med.harvard.edu/dpinteract/), JASPAR (Anthony Mathelier 2016), RegTransBase (Prokaryotes) (Kazakov et al. 2007), Prodoric Release (Prokaryotes) (Munch et al. 2003), and Yeastract (Yeast) (Teixeira et al. 2014), were utilized for regulon filtering in known TF templates to find homologous TFs and corresponding genes in MG1363 using BLAST with default parameters. In the literature validation part, all ASR-related transporters and genes were collected from published articles, and their sequences were obtained from NCBI and UniProt databases.

### Operon identification

The genome-scale operons of MG1363 were identified by DOOR2. It is a one-stop operon-centered resource including operons, alternative transcriptional units, motifs, terminators, and conserved operons information across multiple species (Mao, Ma, Zhou, Chen, Zhang, Yang, Mao, Lai and Xu 2014). Operons were predicted by the back-end prediction algorithm with a prediction accuracy of 90-95%, based on the features of intergenic distance, neighborhood conservation, short DNA motifs, length ratio between gene pairs, and newly developed transcriptomic features trained from the strand-specific RNA-seq dataset (Chou et al. 2015, Li, Liu, et al. 2011).

### Gene differential expression analysis and co-expression analysis

DEGs were identified based on the Wilcoxon signed-rank test (Bauer 1972) between the control and treatment, which was performed in R. The gene co-expression analysis was performed using a hierarchical clustering method (‘hcluster’ package in R) (Antoine Lucas 2006) to detect the CEMs under the acid stress in MG1363.

### Motif finding and regulon prediction

Genes from each CEM were first mapped to the identified operons to retrieve the basic transcription units. Next, 300 bps in the upstream of the translation starting sites for each operon were extracted, in which motif finding was carried out using the web server DMINDA (Ma, Zhang, et al. 2014, Yang et al. 2017). DMINDA is a dominant motif prediction tool, embraced five analytical algorithms to find, scan, and compare motifs (Li, Liu, Ma and Xu 2011, Liu et al. 2017, Ma, Liu, Zhou, Yin, Li and Xu 2013), including a phylogenetic footprint framework to elucidate the mechanism of transcriptional regulation at a system level in prokaryotic genomes (Li, Ma, Mao, Yin, Zhu and Xu 2011, Liu, Zhang, Zhou, Li, Fennell, Wang, Kang, Liu and Ma 2016, Liu, Zhou, Li, Zhang, Zeng, Liu and Ma 2016). All sequences were uploaded to the server and default parameters were used to find the top five significant motifs (*p*-value < 0.05) in each cluster. The identified motifs were subjected to motif comparison and grouped into regulons based a sequence similarity cutoff using the BBC program in DMINDA (Ma, Liu, Zhou, Yin, Li and Xu 2013).

### Regulon validation based on TF BLAST and DEG filtering

Each highly conserved motif was considered to contain the same TFBS among species. Therefore, a comparison study was performed using TOMTOM in the MEME Suite (Bailey et al. 2009) between identified motif and public-domain TFBS databases, including DPINTERACT, JASPAR, RegTransBase, Prodoric Release and Yeastract, to find TFBSs and corresponding TFs with significant *p*-values in other prokaryotic species. Those TFs were then mapped to MG1363 using BLAST by default parameters to predict the connection between regulons and TFs in MG1363. On the other hand, since genes without differential expression were supposed not to react to pH changes, and thus, irrelevant to ASR, regulons without DEGs were not involved in the GRN, and thus, excluded from the following steps.

### Regulon validation based on known ASR proteins from the literature

To validate the performance of the above computational pipeline for regulon prediction, a literature-based validation was performed. Thirty-six ASR-related proteins and genes in other organisms including *L. lactis, E. coli, Streptococci, etc*. were first manually collected from literature, and their sequence was retrieved from the NCBI and UniProt databases. They were used to examine the existing known mechanisms in response to pH changes in MG1363 using the BLAST program by default parameters on NCBI. Such literature-based validation can either confirm the putative regulons when known ASR-related genes can be found in the significant regulons or expand our results to some insufficiently significant regulons, which indicate both false positive and true negative rate to evaluate the computational pipeline.

## RESULTS

### Predicted operons and CEM generation

A total of 1,565 operons with 2,439 coding genes of MG1363 (Dataset S1) were retrieved from the DOOR2 database. Through co-expression analysis, the 1,565 operons were grouped into 124 co-expressed clusters. Among these clusters, two large groupings contain more than 200 operons. Each of which was removed from the subsequent analyses as larger clusters may have higher chances to induce false positive operons which were connected with true operons by co-expression analysis. For the remaining 122 clusters covering 2,122 genes, 26 (21%) contain no more than 10 operons; the smallest cluster had two operons, and most of the clusters (90%) contained between 10 and 50 operons (Dataset S2 and Figure S1).

### Predicted regulons based on motif finding and clustering

Using BoBro in the DMINDA web server, multiple motif sequences were identified from the 300 bps in the upstream of the translation start sites for each operon. Only the top five significant motifs (*p*-value < 0.05) were selected in each cluster, giving rise to a total of 610 (122×5) identified. The motif comparison-and-clustering analysis was then performed on the 610 motifs, and 51 motif clusters were identified with a motif similarity 0.8 as a cutoff. Intuitively, the operons sharing highly similar motifs in each motif cluster are supposed to be regulated by the same TF and tend to be in the same regulon. Hence, these 51 motif clusters correspond to 51 regulons (Dataset S3).

### Computationally-verified regulon based on TF BLAST and differential gene expression (DGE) analysis

Among the above 51 regulons, 14 were found containing motifs significantly (E-value < 0.05) matched to known TFBSs using TOMTOM in the MEME suite, representatively. The motif logos are shown in Figure S2, and more details can be found in Dataset S4. The 14 TFBS-corresponding TFs were then mapped to MG1363 using BLAST to identify the real TFs/genes regulating each regulon. As a result, eight known TFs, spo0A, lhfB, GAL80, CovR, c4494, ihfA, CovR, and RHE_PF00288, were successfully mapped to MG1363 resulting in eight TFs with duplicates. *llrA* (llmg_0908) regulates regulons #12 and #37, *ccpA* (llmg_0775) regulated regulons #15 and #47, *hllA* (llmg_0496) regulates regulons #7 and #31 (Table 1). *CcpA* (Abranches et al. 2008, Zomer et al. 2007), *llrA* (O’Connell-Motherway et al. 2000), *llrC* (O’Connell-Motherway, van Sinderen, Morel-Deville, Fitzgerald, Ehrlich and Morel 2000), and *hllA* (Bolotin et al. 1999) were known to be ASR-related genes in *L. lactis*; gene (llmg_0271), without any related known TF, was found similar to template TF GAL80 in yeast, which has not been associated with any ASR regulation pathways yet. For all 14 significant regulons, regulons #3, #4, #20, #28, #40, and #44 are potential candidates as, currently, no related TFs in *L. lactis* have been found.

**Table 1.** 14 significant regulons that are verified and mapped to known TFs. According to analyses, operon numbers and DEG determination (yes or no), matched template TFs and mapped TFs were assigned for each significant regulon, respectively, and were aligned based on regulon ID number. Five regulons containing DEGs and having the corresponding TF at the same time were bolded, as which were being computationally verified regulons to be responsible for acid stress in MG1363.

| Regulon ID | No. of operons | DEG | TF template | TF(gene) blast in MG1363 |
| --- | --- | --- | --- | --- |
| **Regulon2** | **82** | **Y** | **spo0A** | ***llrC*(llmg_0414)** |
| **Regulon3** | 32 | Y | FoxQ1 | N/A |
| **Regulon4** | 20 | Y | SPT2 | N/A |
| **Regulon7** | **49** | **Y** | **lhfB** | ***hllA*(llmg_0496)** |
| **Regulon10** | 5 | N | GAL80 | -(llmg_0271) |
| **Regulon12** | **259** | **Y** | **CovR** | ***llrA*(llmg_0908)** |
| **Regulon15** | **19** | **Y** | **c4494** | ***ccpA*(llmg_0775)** |
| **Regulon20** | 79 | Y | NHP6A | N/A |
| **Regulon28** | 5 | Y | 1Z916 | N/A |
| **Regulon31** | **65** | **Y** | **ihfA** | ***hllA*(llmg_0496)** |
| **Regulon37** | 10 | N | CovR | *llrA*(llmg_0908) |
| **Regulon40** | 7 | Y | Awh | N/A |
| **Regulon44** | 12 | N | YBR182C | N/A |
| **Regulon47** | 5 | N | RHE_PF00288 | *ccpA*(llmg_0775) |

Additionally, 86 down-regulated genes and 55 up-regulated genes (Dataset S5), resulting from DGE analysis were integrated into the regulons. Regulons #10, #37, #44 and #47 were found lacking DEGs. Thus, gene llmg_0271, related to regulon #10, was not likely to respond to acid stress in MG1363 even though it has been successfully mapped to MG1363, and was then grouped into the potential candidate. On the contrary, *ccpA* and *llrA* were still retained due to their involvements in regulons #15 and #12 with DEGs, respectively.

By the end of the computational pipeline, we predicted that regulons #2, #7, #12, #15 and #31 were related to GRN in MG1363. Merging regulon #7 and #31 as one, we referred their TF names (*ccpA, llrA, llrC*, and *hllA*) to represent the five regulons for convenience.

### Verified regulons based on literature verification

Altogether, 36 literature-supported ASR-related transporters were successfully mapped to MG1363 using blast with an E-value cutoff as 1e-10 and resulted in a total of 33 mapped genes. All the 36 transporters were categorized into nine modules based on their biological functions or regulated pathways, including L-lactate dehydrogenase (LDH), GAD, ADI, urea degradation, F1/F0ATPase, acid stress, protein repair and protease, envelope alterations and DNA repair. The 33 mapped genes generate 22 operons and six regulons: *llrA, llrC, hllA*, NHP6A, regulon #8 and #39, which were subjected, one or more, to each functional module (Table 2).

Regulon *llrA, llrC*, and *hllA* have already been computationally identified in Table 1 and supported again by literature verification results. NHP6A which interestingly has a homologous TF in human and fungi but not in *L. lactis* (Kolodrubetz and Burgum 1990, Stillman 2010), yet failed to map in MG1363. Here, we are using NHP6A to represent regulon #20, as their relationship has been predicted computationally in Table 1. Regulon #39 was identified to be regulated by *llrD*, one of the six two-component regulatory systems in MG1363 (O’Connell-Motherway, van Sinderen, Morel-Deville, Fitzgerald, Ehrlich and Morel 2000). Regulons #8 (llmg_1803) and #39 (*llrD*) were not included in the 14 significant regulons in Table 1. For NHP6A, regulons #8 and #39 were enriched by literature validation as it expanded regulon results of the RECTA pipeline. Among the nine functional modules, *llrA* was found connected to five of them, and NHP6A related to three. On the other hand, the GAD and urea degradation functional modules failed to connect to any previous regulons.

**Table 2.**
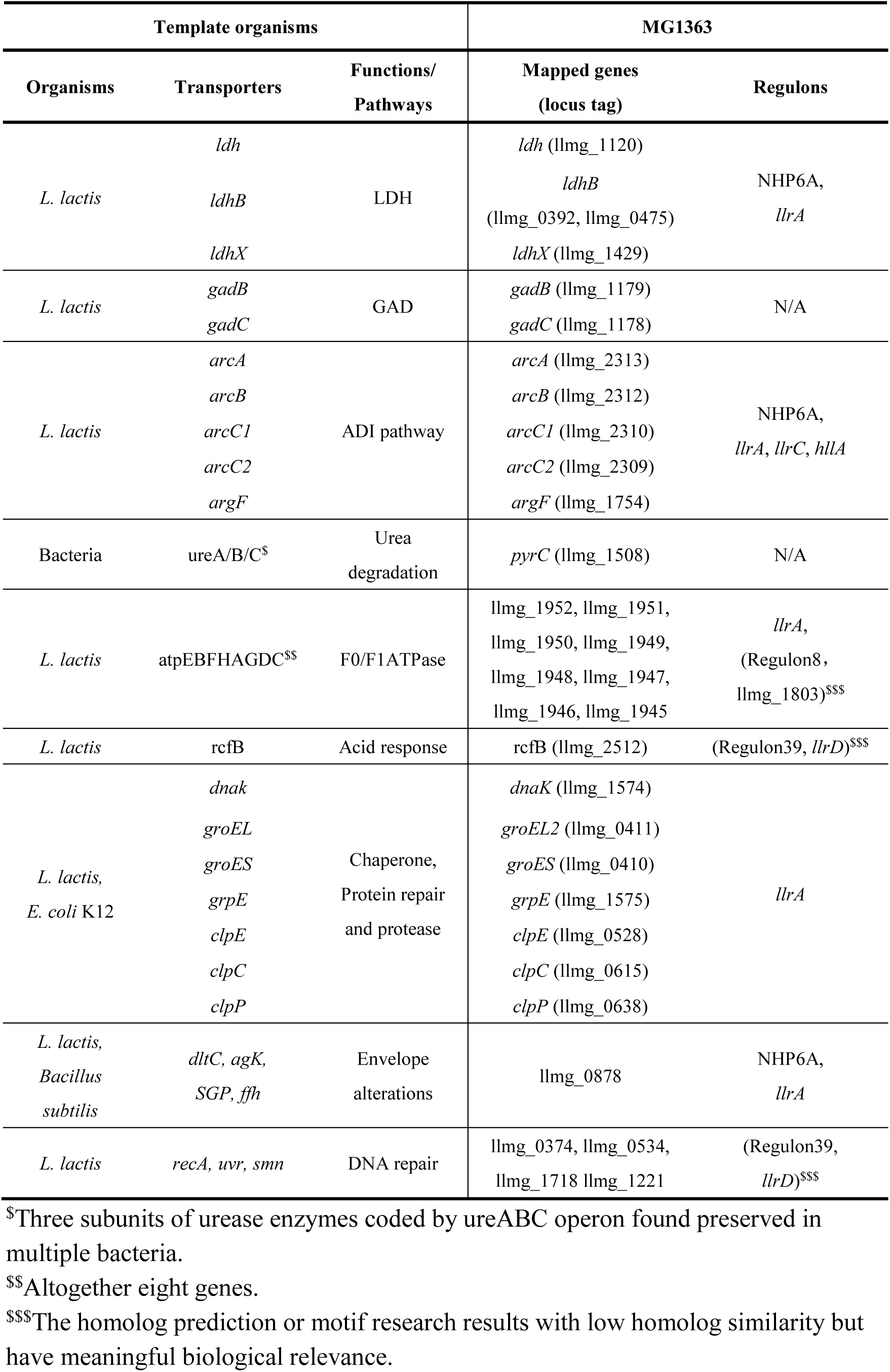
Known ASR-related gene mapping from literature in response to pH change. Literature-supported ASR-related genes found in close species or other *L. lactis* strains. The template transporters and genes were first identified in published studies from the NCBI and UniProt databases. *L. lactis Il1403* was used as the organism which is very close to MG1363 if template gene existed. Only 36 templates that successfully mapped to the MG1363 genome were listed, which resulted in 33 genes. All mapped genes and corresponding templated were organized by their regulated pathways which were further used as functional modules. Mapped genes were searched in 51 regulons to build the connections between functional modules and regulons.

Compared to the regulon verification based on TF BLAST and DGE, the literature verification identified two more regulons (#8 and #39) that lay in the insignificant group, however, with no sign of *ccpA* regulon. Thus, such a result indicates a possible false positive rate of 1/5 and a true negative rate of 2/37 of our computational pipeline, indicating a reliability and feasibility of using RECTA to predict the ASR-related regulons. In Figure 2, we show the processes and results for both literature verification and computational pipeline in detail. The final eight regulons predicted from both parts were then compared to construct a GRN response to acid stress, integrated with other information found in the literature.

**Figure 2.**
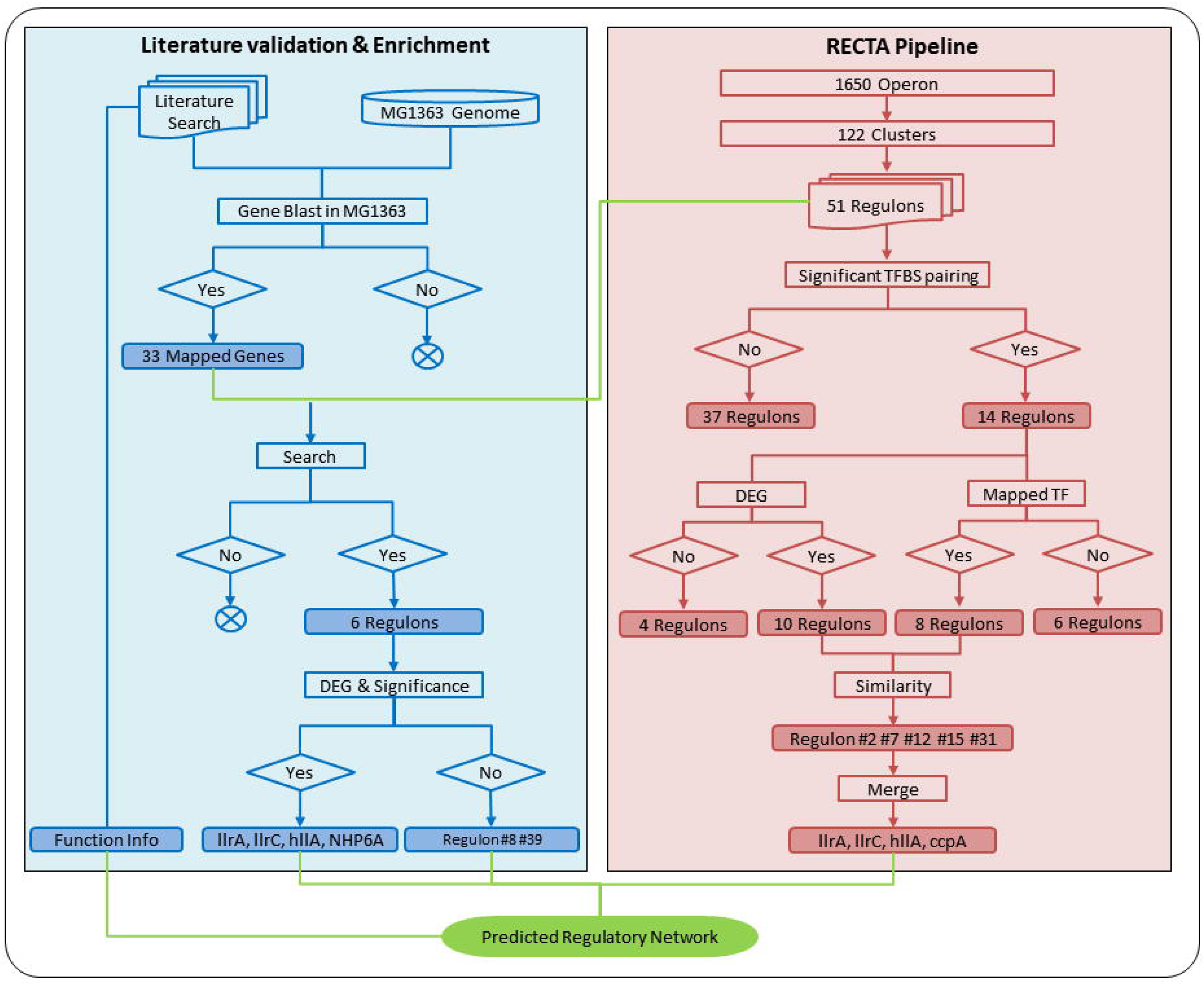
Regulon prediction using RECTA pipeline (red) and validation and enrichment using literature information and gene blast (blue). All processes were shown in rectangles and results were highlighted with corresponding background colors. In the computational pipeline, 51 regulons with assigned motifs and operons were analyzed sequentially through significant TFBS pairing, DEG conformation, and TF blast. Only regulons contained DEGs (ten) and had related mapped TF (eight) were believed to be the final predicted ASR-related regulons (five). These five regulons were then merged into four and using the corresponding TFs to represent their names. In the literature validation process, known ASR-related transporters were first mapped to the MG1363 genome and resulted in 33 genes. Those genes were then searched in 51 regulons and determine six related regulons. All regulons resulted from both computational pipeline and literature validation were combined, along with the information of functional modules, to determine the GRN.

### A model of regulatory network in response to pH change

According to the results outlined above, we are presenting a working model of the transcriptional regulatory network for acid stress response in MG1363 (Figure 3). The network consists of two transmembrane proteins (Dataset S6), eight regulons, nine functional modules, and 33 orthologous genes known for ASR in other bacteria that are also contributing in MG1363.

**Figure 3.**
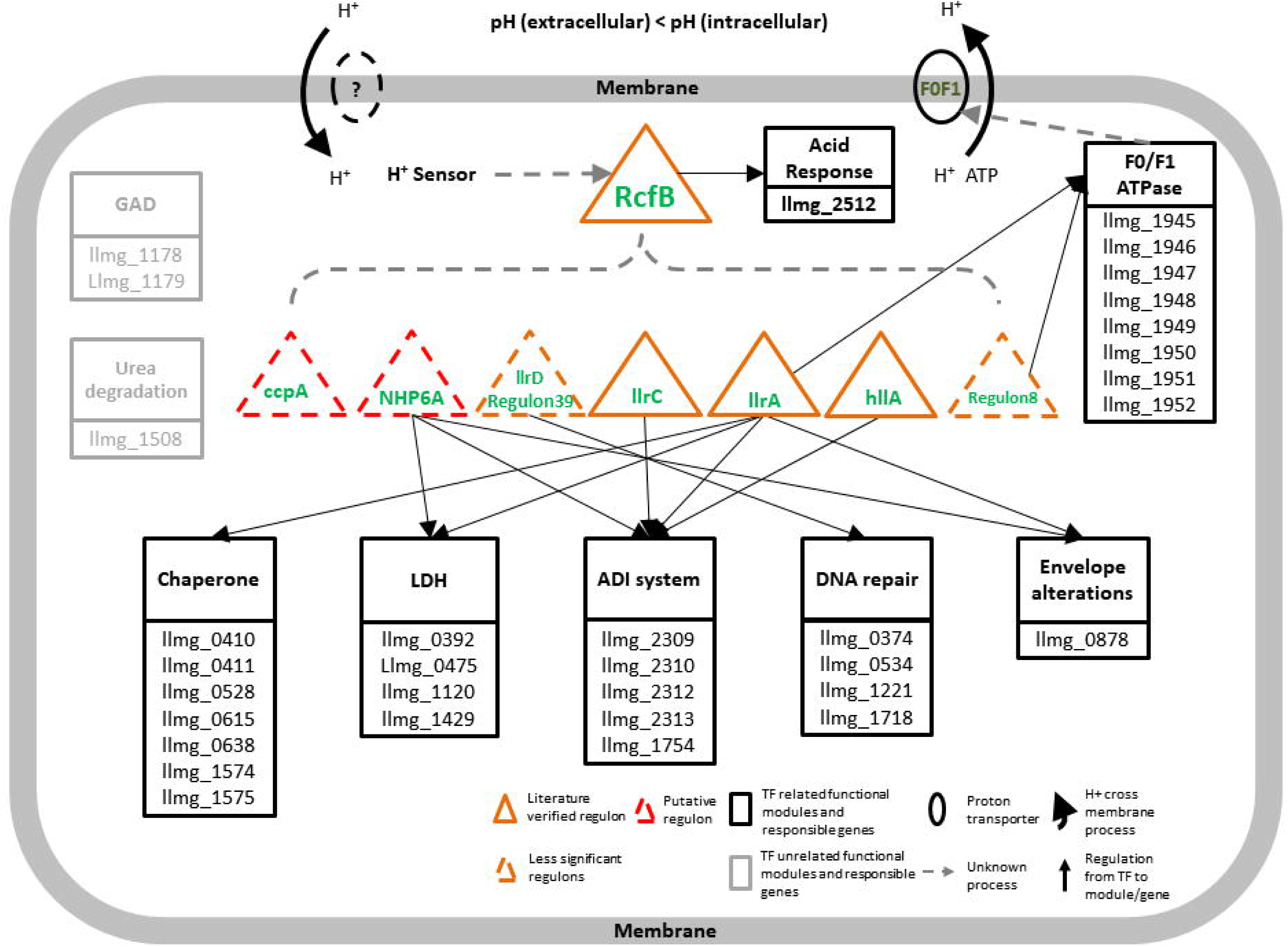
A working model of the transcriptional gene regulatory network in response to pH change in *L. lactis*. The mechanism is activated by the change of proton signal in a cell. *RcfB* is assumed to be the overall activator for the rest seven regulons and controls the ASR functional module solely. Three kinds of literature verified and significant ASR-related regulons, *llrA, llrC* and *hllA*, two insufficient significant regulon *llrD* (regulon #39) and regulon #8 (llmg_1803) predicted via our workflow but results under 0.8 motif similarity cutoff or could not find hit, and one putative significant regulon NHP6A control the seven functional modules which are experimentally verified in close species MG1363. The other significant regulon *ccpA* failed to be confirmed by any literature proved genes or transporters. Two extra functional modules, GAD, and urea degradation show no direct connection to all seven the regulons. One or more homology genes are found in MG1363 for all the nine modules using BLAST. The solid arrows indicate regulation between regulons/TFs and functional modules/genes, and the dashed arrows indicate uncertain control processes. Besides, two ovals indicate two trans-membrane proteins, one isconfirmed as F0/F1ATPase and the other one with dash line that we still not find its related information in the public-domain literature.

The network is subjected to respond to the change of intracellular proton level. The signal is captured by H+ sensor and regulons are initiated to be regulated. Although significance was not shown for *rcfB* in our computational results, it has been reported to recognize and regulate promoter P170 (Madsen et al. 1999), P1 and P3 (Hindre et al. 2004, Rince et al. 1994), which are activated by the ACiD box and essential to acid response (Madsen et al. 2005). With the ACiD box, operons like *groESL, lacticin 481* and *lacZ* have been proved to be regulated by *rcfB*, while *als, aldB*, etc. have not (Madsen, Hindre, Le Pennec, Israelsen and Dufour 2005). The homologous comparative study also predicted the existence of the ACiD box in *llrA* (Akyol et al. 2008, O’Connell-Motherway, van Sinderen, Morel-Deville, Fitzgerald, Ehrlich and Morel 2000). With such evidence, we separated *rcfB* from regulon #39 and predicted that *rcfB* is first triggered by H+ sensor and acts as the global initiator that controls the other seven regulons. It is reasonable that *rcfB*-related regulon #39 failed to show significant TF matching results after CEM treatment in the operon clustering step. *rcfB* worked as a trustworthy global factor; its differential expression should be less significant than regulons directly responding to acid stress, thus leading to the failure of being predicted by the RECTA pipeline. Nevertheless, the low number of microarray data sets (eight) also limited the real performance to the ASR. However, the mechanism of how H+ sensor is activating and regulating the GRN and *rcfB* remains unclear. In the seven regulons, three--*llrA, llrC* and *hllA*--were verified through literature to be related to ASR; regulons #8 and #39 showed less significant in regulon prediction; NHP6A was considered as putative regulon due to its failure to map in MG1363; and *ccpA* was another putative regulon without literature support.

The six downstream regulons (*llrA, llrC, hllA*, NHP6A, regulon #39, and regulon #8) other than *ccpA*, interact with each other to regulate six ASR-related functional modules, including ADI system, DNA repair, LDH, protein repair, envelope alterations, and F0/F1ATPase. The ADI pathway, which generates ATP and protects cells from acid stress (Zuniga, Perez and Gonzalez-Candelas 2002), is under the regulation of NHP6A, *llrC, llrA*, and *hllA*. Another important pathway is the LDH (EC 1.1.1.27) under the regulation of NHP6A and *llrA*, which converts pyruvate and H+ to lactate which is exported outside of cells (Dennis and Kaplan 1960). Chaperons which take part in macromolecule protection and repairing are subjected to regulon *llrA*. Chaperons have functions that includes providing protection to against environmental stress, helping protein folding, and repairing damaged proteins, and have been demonstrated to show clear linkage with acid stress in numerous Gram-positive bacteria (Frees, Vogensen and Ingmer 2003, Jayaraman, Penders and Burne 1997, Kern, Malki, Abdallah, Tagourti and Richarme 2007, Mujacic and Baneyx 2007). F0/F1ATPase, controlled by *llrA* and regulon #8, also plays an important role in maintaining normal cellular pH, which pumps H+ out of cells at the expense of ATP (Amachi et al. 1998, Koebmann, Nilsson, Kuipers and Jensen 2000, Lund, Tramonti and De Biase 2014, O’Sullivan and Condon 1999). The GAD (Nomura, Nakajima, Fujita, Kobayashi, Kimoto, Suzuki and Aso 1999, Sanders, Leenhouts, Burghoorn, Brands, Venema and Kok 1998) and urea degradation (Cotter and Hill 2003) functional modules are missing reliable associations with the regulons in MG1363 while maintaining functions in ASR mechanism in other species.

## DISCUSSION AND CONCLUSIONS

Implementation of the novel computational pipeline RECTA resulted in the construction of an eight-regulons enrolled ASR regulatory network. The framework provides a useful tool and will be a starting point toward a more systems-level understanding of the question (Cao, Wei, et al. 2017). The identified motifs and regulons suggest acid resistance is a coordinated response regarding regulons, although most of which have not been identified or experimentally verified. From the three well-identified regulons—*llrA, llrC*, and *hllA*, it appears the gene regulation is also complex, as these regulons also interact with other proteins and TFs. F0/F1ATPase is directly involved in the concentration regulations of the intracellular proton. Other pathways are responsible for repairing the damage caused by acid stress, such as DNA repair, protein repair, and cell envelops alterations. However, there were also several reported ASR-related genes or transporters such as *htrA* in *Clostridium spp*. (Alsaker et al. 2010), CovS/CovR acid response regulator in *Streptococcus* (Cumley et al. 2012), cyclopropane fatty acid (cfa) synthase for cell-membrane modification (Budin-Verneuil et al. 2005), and oxidative damage protectant genes like *sodA, nox-1* and *nox-2* (Santi et al. 2009) that failed to map to MG1363. Using more gene expression datasets for CEM and DGE analyses could be a way to strengthen the result of our computational pipeline, which might cover more significant regulons to construct a more solid and complete regulatory network.

Homology mapping at the genomic level showed very a long evolutionary distance between MG1363 and currently well-annotated model species. Hence, the functional analysis for MG1363 is limited, and it is hard to apply gene functional enrichment to verify our prediction results. With more expression datasets and experiments about protein-protein interactions, the ASR mechanism can be largely improved in *L. lactic* MG1363.

In summary, through the implementation of RECTA, we found that the ASR at the transcriptome level in MG1363 is an orchestrated complex network. Functional annotation shows these regulons are involved in many levels of biological processes, including but not limited to DNA expression, transcription, and metabolism. Our method builds a TF-regulons-GRN relationship so that the new ASR-related genes can be predicted. Besides, the low false positive and true negative rate indicate the RECTA pipeline as sensitive and reasonable. In fact, considering the high accuracy, we regarded *ccpA* as the putative regulon, though not connected to any related functional modules, while more robust methods are required. Such results expand current pathways to those that can corroborate cell structures—cell wall, cell membranes, etc.—and related functions. Our findings suggest that acid has profound adverse effects and inflict systems-level response. Such predicted response pathways can inform better resistance design.

Looking forward to the acid tolerance advantage of *L. lactis*, which makes its prospective application in drug and vaccine delivery, the effects on anti-obesity research, and metagenomic studies, the ASR-related GRN in *L. lactis* shows an excellent research value. Fully understanding of its theory may contribute to the development of *Lactococcus* therapy and can even expand to other close species by genetic modification. Furthermore, our computational pipeline provides an effective method to construct reliable GRN based on regulon prediction, integrating CEMs, DGE analysis, motif finding, and comparative genomics study. It has a durable application power and can be effectively applied to other bacterial genomes, where the elucidation of the transcriptional regulation network is needed.

In this study, we designed a computational framework, RECTA, for acid-response regulon elucidation. This tool integrates differential gene expression, co-expression analysis, *cis-*regulatory motif identification, and comparative genomics to predict and validate regulons associated with acid response. In demonstrating the efficacy of this tool, we analyzed *Lactococcus lactis* MG1363. This implementation resulted in the expanded understanding of the acid-response regulon network for this one strain of *L. lactis* and provides an applicable method for acid-response regulon elucidation of further species. Through utilization of the RECTA pipeline, researchers can readily evaluate acid-response mechanisms for numerous bacterial species, while simultaneously validating the results of their study.

## LIST OF ABBREVIATIONS

ADI: arginine deiminase
(BBC): BoBro-based motif Comparison
(BoBro): BOttleneck BROken
CEM: co-expression (gene) module
DEG: differentially expressed gene
DGE: differential gene expression
*E. coli*: *Escherichia coli*
GAD: glutamate decarboxylases
GRN: gene regulatory network
LDH: lactate dehydrogenase
*L. lactis*: *Lactococcus lactis*
MG1363: *Lactococcus lactis* MG1363
Motif: cis-regulatory motif
RECTA: regulon identification based on comparative genomics and transcriptomics analysis
TF: transcription factors
TFBS: transcriptional factor binding site

## DECLARATIONS

### Ethics Approval and Consent to Participate

Not applicable

### Data Availability Statements

All datasets analyzed and generated for this study are included in the manuscript and the supplementary files.

### Authors’ Contributions

XC and QM carried out the framework of the paper and are responsible for the regulon prediction work. HZ carried out the operon prediction and motif finding work. CL was responsible for DGE analysis. HC, A. McDermaid, and A. Ma drafted and revised the manuscript. All authors submitted comments, read and approved the final manuscript.

## SUPPLEMENTAL MATERIALS

**Supplement Number 1:** Figure S1 operon_distribution.pptx; The distribution of operon numbers among 122 clusters.

**Supplement Number 2:** Figure S2 motif_logos_for_14_significant_regulons.pptx; The motif logos substracted from 14 significant regulons, used for TFBS comparison.

**Supplement Number 3:** Dataset S1 NC_009004_operons_and_genes.xlsx; The operon prediction results for MG1363.

**Supplement Number 4:** Dataset S2 122_operons_to_clusters.xlsx; The gene clustering results for microarray “GSE47012” containing operons.

**Supplement Number 5:** Dataset S3 51_motif-regulons.xlsx; The summarized motif prediction results for regulon prediction.

**Supplement Number 6:** Dataset S4 14_verified_regulons.xlsx; Computational validation results for predicted regulons.

**Supplement Number 7:** Dataset S5 differential_expression_genes.xlsx; 86 up-regulated genes and 55 down-regulated genes.

**Supplement Number 8:** Dataset S6 trans-memberane.xlsx; The prediction results for trans-membrane proteins.

## REFERENCES

Abranches J, Nascimento MM, Zeng L, Browngardt CM, Wen ZT, Rivera MF, Burne RA. 2008. CcpA regulates central metabolism and virulence gene expression in Streptococcus mutans. J Bacteriol. Apr;190:2340–2349. Epub 2008/01/29.

Aibar S, Gonzalez-Blas CB, Moerman T, Huynh-Thu VA, Imrichova H, Hulselmans G, Rambow F, Marine JC, Geurts P, Aerts J, et al 2017. SCENIC: single-cell regulatory network inference and clustering. Nat Methods. Nov;14:1083–1086. Epub 2017/10/11.

Akyol I, Comlekcioglu U, Karakas A, Serdaroglu K, Ekinci MS, Ozkose E. 2008. Regulation of the acid inducible rcfB promoter in Lactococcus lactis subsp. lactis. Annals of Microbiology. 2008//;58:269.

Alsaker KV, Paredes C, Papoutsakis ET. 2010. Metabolite stress and tolerance in the production of biofuels and chemicals: gene-expression-based systems analysis of butanol, butyrate, and acetate stresses in the anaerobe Clostridium acetobutylicum. Biotechnol Bioeng. Apr 15;105:1131–1147. Epub 2009/12/10.

Amachi S, Ishikawa K, Toyoda S, Kagawa Y, Yokota A, Tomita F. 1998. Characterization of a mutant of Lactococcus lactis with reduced membrane-bound ATPase activity under acidic conditions. Biosci Biotechnol Biochem. Aug;62:1574–1580. Epub 1998/10/03.

Anthony Mathelier OF, David J. Arenillas, Chih-yu Chen, Grégoire Denay, Jessica Lee, Wenqiang Shi, Casper Shyr, Ge Tan, Rebecca Worsley-Hunt Allen W. Zhang, François Parcy, Boris Lenhard, Albin Sandelin, Wyeth W. Wasserman. 2016. JASPAR 2016: a major expansion and update of the open-access database of transcription factor binding profiles Nucleic Acids Research. 03 November 2015;44:D110–D115.

Antoine Lucas SJ. 2006. Using amap and ctc Packages for Huge Clustering. R News.6:58–60.

Arnoldini M, Mostowy R, Bonhoeffer S, Ackermann M. 2012. Evolution of Stress Response in the Face of Unreliable Environmental Signals. PLoS Comput Biol.8:e1002627.

Bailey TL, Boden M, Buske FA, Frith M, Grant CE, Clementi L, Ren J, Li WW, Noble WS. 2009. MEME Suite: tools for motif discovery and searching. Nucleic Acids Research. July 1, 2009;37:W202–W208.

Bauer DF. 1972. Constructing Confidence Sets Using Rank Statistics. Journal of the American Statistical Association.67:687–690.

Bermudez-Humaran LG, Cortes-Perez NG, Lefevre F, Guimaraes V, Rabot S, Alcocer-Gonzalez JM, Gratadoux JJ, Rodriguez-Padilla C, Tamez-Guerra RS, Corthier G, et al 2005. A novel mucosal vaccine based on live Lactococci expressing E7 antigen and IL-12 induces systemic and mucosal immune responses and protects mice against human papillomavirus type 16-induced tumors. J Immunol. Dec 01;175:7297–7302. Epub 2005/11/23.

Blanchette M, Schwikowski B, Tompa M. 2002. Algorithms for phylogenetic footprinting. J Comput Biol. 9:211–223. Epub 2002/05/23.

Bolotin A, Mauger S, Malarme K, Ehrlich SD, Sorokin A. 1999. Low-redundancy sequencing of the entire Lactococcus lactis IL1403 genome. Antonie Van Leeuwenhoek. Jul-Nov;76:27–76. Epub 1999/10/26.

Budin-Verneuil A, Maguin E, Auffray Y, Ehrlich DS, Pichereau V. 2006. Genetic structure and transcriptional analysis of the arginine deiminase (ADI) cluster in Lactococcus lactis MG1363. Can J Microbiol. Jul;52:617–622.

Budin-Verneuil A, Maguin E, Auffray Y, Ehrlich SD, Pichereau V. 2005. Transcriptional analysis of the cyclopropane fatty acid synthase gene of Lactococcus lactis MG1363 at low pH. FEMS Microbiol Lett. Sep 15;250:189–194. Epub 2005/08/16.

Cao H, Ma Q, Chen X, Xu Y. 2017. DOOR: a prokaryotic operon database for genome analyses and functional inference. Brief Bioinform. Jul 28. Epub 2017/10/03.

Cao H, Wei D, Yang Y, Shang Y, Li G, Zhou Y, Ma Q, Xu Y. 2017. Systems-level understanding of ethanol-induced stresses and adaptation in E. coli. Scientific Reports. 03/16/online;7:44150.

Carvalho AL, Turner DL, Fonseca LL, Solopova A, Catarino T, Kuipers OP, Voit EO, Neves AR, Santos H. 2013. Metabolic and transcriptional analysis of acid stress in Lactococcus lactis, with a focus on the kinetics of lactic acid pools. PLOS ONE.8:e68470.

Chou WC, Ma Q, Yang S, Cao S, Klingeman DM, Brown SD, Xu Y. 2015. Analysis of strand-specific RNA-seq data using machine learning reveals the structures of transcription units in Clostridium thermocellum. Nucleic Acids Res. May 26;43:e67. Epub 2015/03/15.

Cotter PD, Hill C. 2003. Surviving the acid test: responses of gram-positive bacteria to low pH. Microbiol Mol Biol Rev. Sep;67:429–453, table of contents. Epub 2003/09/11.

Cumley NJ, Smith LM, Anthony M, May RC. 2012. The CovS/CovR acid response regulator is required for intracellular survival of group B Streptococcus in macrophages. Infect Immun. May;80:1650–1661. Epub 2012/02/15.

Dennis D, Kaplan NO. 1960. D-and L-lactic acid dehydrogenases in Lactobacillus plantarum. J Biol Chem. Mar;235:810–818. Epub 1960/03/01.

Frees D, Vogensen FK, Ingmer H. 2003. Identification of proteins induced at low pH in Lactococcus lactis. International Journal of Food Microbiology. 11/1/;87:293–300.

Gupta S, Stamatoyannopoulos JA, Bailey TL, Noble WS. 2007. Quantifying similarity between motifs. Genome Biol.8:R24. Epub 2007/02/28.

Hanniffy SB, Carter AT, Hitchin E, Wells JM. 2007. Mucosal delivery of a pneumococcal vaccine using Lactococcus lactis affords protection against respiratory infection. J Infect Dis. Jan 15;195:185–193. Epub 2006/12/28.

Hartke A, Bouché S, Giard J-C, Benachour A, Boutibonnes P, Auffray Y. 1996. The lactic acid stress response of Lactococcus lactis subsp. lactis. Current Microbiology.33:194–199.

Hauryliuk V, Atkinson GC, Murakami KS, Tenson T, Gerdes K. 2015. Recent functional insights into the role of (p)ppGpp in bacterial physiology. Nat Rev Micro. 05//print;13:298–309.

Hindre T, Le Pennec JP, Haras D, Dufour A. 2004. Regulation of lantibiotic lacticin 481 production at the transcriptional level by acid pH. FEMS Microbiol Lett. Feb 16;231:291–298. Epub 2004/02/28.

Hols P, Kleerebezem M, Schanck AN, Ferain T, Hugenholtz J, Delcour J, de Vos WM. 1999. Conversion of Lactococcus lactis from homolactic to homoalanine fermentation through metabolic engineering. Nat Biotechnol. Jun;17:588–592. Epub 1999/06/29.

Hoskins J, Alborn WE, Jr., Arnold J, Blaszczak LC, Burgett S, DeHoff BS, Estrem ST, Fritz L, Fu DJ, Fuller W, et al 2001. Genome of the bacterium Streptococcus pneumoniae strain R6. J Bacteriol. Oct;183:5709–5717. Epub 2001/09/07.

Hutkins RW, Nannen NL. 1993. pH homeostasis in lactic acid bacteria. Journal of Dairy Science. 1993/08/01;76:2354–2365.

Jacob F, Perrin D, Sanchez C, Monod J. 1960. [Operon: a group of genes with the expression coordinated by an operator]. C R Hebd Seances Acad Sci. Feb 29;250:1727–1729. Epub 1960/02/29.

Jayaraman GC, Penders JE, Burne RA. 1997. Transcriptional analysis of the Streptococcus mutans hrcA, grpE and dnaK genes and regulation of expression in response to heat shock and environmental acidification. Mol Microbiol. Jul;25:329–341. Epub 1997/07/01.

Jensen ST, Shen L, Liu JS. 2005. Combining phylogenetic motif discovery and motif clustering to predict co-regulated genes. Bioinformatics. Oct 15;21:3832–3839. Epub 2005/08/18.

Katara P, Grover A, Sharma V. 2012. Phylogenetic footprinting: a boost for microbial regulatory genomics. Protoplasma. Oct;249:901–907. Epub 2011/11/25.

Kazakov AE, Cipriano MJ, Novichkov PS, Minovitsky S, Vinogradov DV, Arkin A, Mironov AA, Gelfand MS, Dubchak I. 2007. RegTransBase--a database of regulatory sequences and interactions in a wide range of prokaryotic genomes. Nucleic Acids Res. Jan;35:D407–412. Epub 2006/12/05.

Kern R, Malki A, Abdallah J, Tagourti J, Richarme G. 2007. Escherichia coli HdeB is an acid stress chaperone. J Bacteriol. Jan;189:603–610. Epub 2006/11/07.

Koebmann BJ, Nilsson D, Kuipers OP, Jensen PR. 2000. The membrane-bound H+-ATPase complex is essential for growth of Lactococcus lactis. Journal of Bacteriology. September 1, 2000; 182:4738–4743.

Kolodrubetz D, Burgum A. 1990. Duplicated NHP6 genes of Saccharomyces cerevisiae encode proteins homologous to bovine high mobility group protein 1. J Biol Chem. Feb 25;265:3234–3239. Epub 1990/02/25.

Kumka JE, Bauer CE. 2015. Analysis of the FnrL regulon in Rhodobacter capsulatus reveals limited regulon overlap with orthologues from Rhodobacter sphaeroides and Escherichia coli. BMC Genomics. Nov 04;16:895. Epub 2015/11/06.

Levine JH, Lin Y, Elowitz MB. 2013. Functional roles of pulsing in genetic circuits. Science. 342:1193–1200.

Li G, Liu B, Ma Q, Xu Y. 2011. A new framework for identifying cis-regulatory motifs in prokaryotes. Nucleic Acids Res. Apr;39:e42. Epub 2010/12/15.

Li G, Ma Q, Mao X, Yin Y, Zhu X, Xu Y. 2011. Integration of sequence-similarity and functional association information can overcome intrinsic problems in orthology mapping across bacterial genomes. Nucleic Acids Res. Dec;39:e150. Epub 2011/10/04.

Linares DM, Kok J, Poolman B. 2010. Genome sequences of Lactococcus lactis MG1363 (revised) and NZ9000 and comparative physiological studies. Journal of Bacteriology. November 1, 2010;192:5806–5812.

Liu B, Yang J, Li Y, McDermaid A, Ma Q. 2017. An algorithmic perspective of de novo cis-regulatory motif finding based on ChIP-seq data. Brief Bioinform. Mar 08. Epub 2017/03/24.

Liu B, Zhang H, Zhou C, Li G, Fennell A, Wang G, Kang Y, Liu Q, Ma Q. 2016. An integrative and applicable phylogenetic footprinting framework for cis-regulatory motifs identification in prokaryotic genomes. BMC genomics. Aug 09;17:578. Epub 2016/08/11.

Liu B, Zhou C, Li G, Zhang H, Zeng E, Liu Q, Ma Q. 2016. Bacterial regulon modeling and prediction based on systematic cis regulatory motif analyses. Sci Rep. Mar 15;6:23030. Epub 2016/03/16.

Locke JCW, Young JW, Fontes M, Jiménez MJH, Elowitz MB. 2011. Stochastic pulse regulation in bacterial stress response. Science. 334:366–369.

Lund P, Tramonti A, De Biase D. 2014. Coping with low pH: molecular strategies in neutralophilic bacteria. FEMS Microbiol Rev. Nov;38:1091–1125. Epub 2014/06/06.

Ma Q, Liu B, Zhou C, Yin Y, Li G, Xu Y. 2013. An integrated toolkit for accurate prediction and analysis of cis-regulatory motifs at a genome scale. Bioinformatics. Sep 15;29:2261–2268. Epub 2013/07/13.

Ma Q, Zhang H, Mao X, Zhou C, Liu B, Chen X, Xu Y. 2014. DMINDA: an integrated web server for DNA motif identification and analyses. Nucleic Acids Research. April 21, 2014.

Ma Y, Liu J, Hou J, Dong Y, Lu Y, Jin L, Cao R, Li T, Wu J. 2014. Oral administration of recombinant Lactococcus lactis expressing HSP65 and tandemly repeated P277 reduces the incidence of type I diabetes in non-obese diabetic mice. PLoS One.9:e105701. Epub 2014/08/27.

Madsen SM, Arnau J, Vrang A, Givskov M, Israelsen H. 1999. Molecular characterization of the pH-inducible and growth phase-dependent promoter P170 of Lactococcus lactis. Mol Microbiol. Apr;32:75–87. Epub 1999/04/27.

Madsen SM, Hindre T, Le Pennec JP, Israelsen H, Dufour A. 2005. Two acid-inducible promoters from Lactococcus lactis require the cis-acting ACiD-box and the transcription regulator RcfB. Mol Microbiol. May;56:735–746. Epub 2005/04/12.

Mao X, Ma Q, Liu B, Chen X, Zhang H, Xu Y. 2015. Revisiting operons: an analysis of the landscape of transcriptional units in E. coli. BMC Bioinformatics. Nov 04;16:356. Epub 2015/11/06.

Mao X, Ma Q, Zhou C, Chen X, Zhang H, Yang J, Mao F, Lai W, Xu Y. 2014. DOOR 2.0: presenting operons and their functions through dynamic and integrated views. Nucleic Acids Research. January 1, 2014;42:D654–D659.

Million M, Maraninchi M, Henry M, Armougom F, Richet H, Carrieri P, Valero R, Raccah D, Vialettes B, Raoult D. 2012. Obesity-associated gut microbiota is enriched in Lactobacillus reuteri and depleted in Bifidobacterium animalis and Methanobrevibacter smithii. Int J Obes (Lond). Jun;36:817–825. Epub 2011/08/11.

Mujacic M, Baneyx F. 2007. Chaperone Hsp31 contributes to acid resistance in stationary-phase Escherichia coli. Appl Environ Microbiol. Feb;73:1014–1018. Epub 2006/12/13.

Munch R, Hiller K, Barg H, Heldt D, Linz S, Wingender E, Jahn D. 2003. PRODORIC: prokaryotic database of gene regulation. Nucleic Acids Res. Jan 01;31:266–269. Epub 2003/01/10.

Nomura M, Nakajima I, Fujita Y, Kobayashi M, Kimoto H, Suzuki I, Aso H. 1999. Lactococcus lactis contains only one glutamate decarboxylase gene. Microbiology. Jun;145 (Pt 6):1375–1380.

Novichkov PS, Rodionov DA, Stavrovskaya ED, Novichkova ES, Kazakov AE, Gelfand MS, Arkin AP, Mironov AA, Dubchak I. 2010. RegPredict: an integrated system for regulon inference in prokaryotes by comparative genomics approach. Nucleic Acids Res. Jul;38:W299–307. Epub 2010/06/15.

O’Connell-Motherway M, van Sinderen D, Morel-Deville F, Fitzgerald GF, Ehrlich SD, Morel P. 2000. Six putative two-component regulatory systems isolated from Lactococcus lactis subsp. cremoris MG1363. Microbiology. Apr;146 (Pt 4):935–947. Epub 2000/04/28.

O’Sullivan E, Condon S. 1999. Relationship between acid tolerance, cytoplasmic pH, and ATP and H+- ATPase levels in chemostat cultures of Lactococcus lactis. Appl Environ Microbiol. Jun;65:2287–2293. Epub 1999/05/29.

Rallu F, Gruss A, Ehrlich SD, Maguin E. 2000. Acid-and multistress-resistant mutants of Lactococcus lactis: identification of intracellular stress signals. Molecular Microbiology.35:517–528.

Ramasamy R, Yasawardena S, Zomer A, Venema G, Kok J, Leenhouts K. 2006. Immunogenicity of a malaria parasite antigen displayed by Lactococcus lactis in oral immunisations. Vaccine. May 01;24:3900–3908. Epub 2006/03/21.

Rince A, Dufour A, Le Pogam S, Thuault D, Bourgeois CM, Le Pennec JP. 1994. Cloning, expression, and nucleotide sequence of genes involved in production of lactococcin DR, a bacteriocin from lactococcus lactis subsp. lactis. Appl Environ Microbiol. May;60:1652–1657. Epub 1994/05/01.

Ryan S, Begley M, Gahan CG, Hill C. 2009. Molecular characterization of the arginine deiminase system in Listeria monocytogenes: regulation and role in acid tolerance. Environ Microbiol. Feb;11:432–445. Epub 2009/02/07.

Sanders JW, Leenhouts K, Burghoorn J, Brands JR, Venema G, Kok J. 1998. A chloride-inducible acid resistance mechanism in Lactococcus lactis and its regulation. Mol Microbiol. Jan;27:299–310. Epub 1998/03/04.

Sanders JW, Venema G, Kok J. 1999. Environmental stress responses in Lactococcus lactis. FEMS Microbiology Reviews.23:483–501.

Santi I, Grifantini R, Jiang SM, Brettoni C, Grandi G, Wessels MR, Soriani M. 2009. CsrRS regulates group B Streptococcus virulence gene expression in response to environmental pH: a new perspective on vaccine development. J Bacteriol. Sep;191:5387–5397. Epub 2009/06/23.

Shabayek S, Spellerberg B. 2017. Acid Stress Response Mechanisms of Group B Streptococci. Front Cell Infect Microbiol.7:395. Epub 2017/09/25.

Stillman DJ. 2010. Nhp6:a small but powerful effector of chromatin structure in Saccharomyces cerevisiae. Biochim Biophys Acta. Jan-Feb;1799:175–180. Epub 2010/02/04.

Sun Y, Fukamachi T, Saito H, Kobayashi H. 2012. Adenosine deamination increases the survival under acidic conditions in Escherichia coli. J Appl Microbiol. Apr;112:775–781. Epub 2012/01/27.

Tan K, Moreno-Hagelsieb G, Collado-Vides J, Stormo GD. 2001. A comparative genomics approach to prediction of new members of regulons. Genome Res. Apr;11:566–584. Epub 2001/04/03.

Teixeira MC, Monteiro PT, Guerreiro JF, Goncalves JP, Mira NP, dos Santos SC, Cabrito TR, Palma M, Costa C, Francisco AP, et al 2014. The YEASTRACT database: an upgraded information system for the analysis of gene and genomic transcription regulation in Saccharomyces cerevisiae. Nucleic Acids Res. Jan;42:D161–166. Epub 2013/10/31.

van de Guchte M, Serror P, Chervaux C, Smokvina T, Ehrlich SD, Maguin E. 2002. Stress responses in lactic acid bacteria. Antonie van Leeuwenhoek.82:187–216.

Wegmann U, O’Connell-Motherway M, Zomer A, Buist G, Shearman C, Canchaya C, Ventura M, Goesmann A, Gasson MJ, Kuipers OP, et al 2007. Complete Genome Sequence of the Prototype Lactic Acid Bacterium Lactococcus lactis subsp. cremoris MG1363. Journal of Bacteriology. April 15, 2007;189:3256–3270.

Yang J, Chen X, McDermaid A, Ma Q. 2017. DMINDA 2 0: integrated and systematic views of regulatory DNA motif identification and analyses. Bioinformatics. Apr 13.

Zhang B, Li A, Zuo F, Yu R, Zeng Z, Ma H, Chen S. 2016. Recombinant Lactococcus lactis NZ9000 secretes a bioactive kisspeptin that inhibits proliferation and migration of human colon carcinoma HT-29 cells. Microb Cell Fact. Jun 10;15:102. Epub 2016/06/12.

Zhou C, Ma Q, Li G. 2014. Elucidation of operon structures across closely related bacterial genomes. PLoS One.9:e100999. Epub 2014/06/25.

Zomer AL, Buist G, Larsen R, Kok J, Kuipers OP. 2007. Time-resolved determination of the CcpA regulon of Lactococcus lactis subsp. cremoris MG1363. J Bacteriol. Feb;189:1366–1381. Epub 2006/10/10.

Zuniga M, Perez G, Gonzalez-Candelas F. 2002. Evolution of arginine deiminase (ADI) pathway genes. Mol Phylogenet Evol. Dec;25:429–444. Epub 2002/11/27.

